# A new insight for the screening of potential β-lactamase inhibitors

**DOI:** 10.1101/005181

**Authors:** Vijai Singh, Dharmendra Kumar Chaudhary

## Abstract

The β-lactamase produces by *Aeromonas hydrophila* which enables to hydrolyze and inactivate β- lactam ring of antibiotics. The homology modeling was used to generate the 3-D model of β-lactamase by using known template 3-D structure. The stereochemical quality and torsion angle of 3-D model were validated. Total eleven effective drugs have been selected and targeted the active amino acid residues in β-lactamase. The drugs were derivative of β-lactam ring antibiotics and screening was made by docking. Out of 11 drugs, 3 drugs (Ampicillin, Astreonam and Sultamicillin) were found to be more potent on the basis of robust binding energy between protein-drug interactions. Additionally, homology of β-lactamase of *A. hydrophila* resembled with other pathogenic bacteria that used for phylogeny analysis. These findings suggest a new insight for better understanding and useful for designing of novel potent drugs.

## 1. INTRODUCTION

*Aeromonas hydrophila* is an opportunistic bacterial pathogen which causes hemorrhagic septicemia in fish, reptiles, amphibians (Austin and Austin 1999) and soft tissues infection and diarrhea in human (Barghouthi et al., 1989). There are fifty cases of *Aeromonas* septicaemia reported in severe hepatic cirrhosis and found 52% *A. hydrophila* isolates. A case of liver cirrhosis with *A. hydrophila* infection presenting as acute gastroenteritis and non-traumatic acute osteomyelitis. It has been shown that *A. hydrophila* frequently affects immunocompromised patient with liver cirrhosis (Lee et al., 2003). *A. hydrophila* secretes a number of extracellular enzymes including proteases, DNase, RNase, elastase, lecithinase, amylase, lipase, gelatinase, chitinase (Merino et al., 1995; Pemberton et al., 1997) and cytotoxic/cytolytic enterotoxins (Chopra et al., 1993) and three haemolysins (Hanes and Chandler 1993; Hirono and Aoki 1991; Howard and Buckley 1985). There is a pressing need to diagnose and cure *A. hydrophila* infections.

*A. hydrophila* have been studied for antibiotic sensitivity assay and found all isolates (total no. 25) were resistant to Cephalothin, Ampicillin, Novobiocin and Nitrofurazone, and sensitive to Gentamicin (80%), Co-trimaxazole (92%), Chloramphenicol and Ciprofloxacin (Rathore et al., 2006). An urgent need arises to develop vaccine for controlling of infection but not available yet. Therefore, antibiotics are an alternative and widely used for controlling of *A. hydrophila* infection in human and other animals. β-lactamase is one of the most significant enzymes produces from *A*. *hydrophila*. The β-Lactam antibiotics have been used for controlling of microbial infections. Bacteria have been evolved to hydrolyze the β-lactams by production of β-lactamases (Frere 1995). There is heterogeneity of β-lactamases thus, inhibitors are ineffective. However, some inhibitors are effective only against serine β-lactamases as they are hydrolyzed by metallo β–lactamases (MBLs). A number of inhibitors for MBLs have been reported their side chains binding in a predominantly hydrophobic pocket while their functional groups interacted with zinc ions (Heinz et al., 2003; Siemann et al., 2002).

The 3-D model of β-lactamase of *A. hydrophila* is unknown so for. The homology modeling is used to generate the 3-D model of β-lactamase by using the known 3-D crystal structure as a template. While molecular docking is used for screening of potent and specific drugs by targeting active amino acid residue in β-lactamase. There are a number of previous reports for used of homology modelling of β- ketoacyl acyl carrier protein synthase (KAS) III of *Enterococcus faecalis.* Similarly, homology modeling has been used to generate the 3D model of KASIII protein. The identification of active site residue has been performed using docking and found the two antibacterial drugs for inhibition of growth (Jeong et al., 2007). In this study, homology modeling has also been used for construction of 3-D model of NAD^+^ dependent DNA ligase of *Mycobacterium tuberculosis*. The screening of a number of drugs has been performed by docking approach (Srivastava et al., 2005). The phylogenetic relationship based of protein homology is vital for better understanding of genetic evolutionary relationship of organisms. The aim of present study is to generate the 3-D model of β-lactamase and screening of potent drugs.

## 2. MATERIALS AND METHODS

### 2.1 Retrieval and searching of sequences

The complete protein sequences of β-lactamase from *A. hydrophila* and other bacteria strains were retrieved from National Centre for Biotechnology Information (http://www.ncbi.nlm.nih.gov). The relatedness of sequences deposited in databases was evaluated by BLAST (Basic Local Alignment Search Tool) (Altschul et al., 1990) that was implemented via NCBI (http://www.ncbi.nlm.nih.gov/blast). The BlastP (protein query–protein database comparison) was performed with protein data bank (PDB). The alignment was also performed with target protein sequences with template protein (PDB: 1X8G) using CLUSTAL X 1.83.

### 2.2 Generation and evaluation of 3-D model

The X-crystal structural mono zinc carbapenemase (Cpha) from *A. hydrophila* was available at 1.70 Å resolution (PDB: 1X8G) that was used as template structure to generate 3-D model of β-lactamase. The homology modeling was used to generate the 3D structure of β-lactamase through the Modeller9v2. It was evaluated by minimum model score and dope score of model and template. The 3D model was further validated by PROCHECK that also created a Ramachandran plot. The accuracy and overall G-factor were also considered for model evaluation.

### 2.3 Screening of potent drugs

The drugs were taken from NCBI Pubchem compounds in 2-D structure that was converted into 3-D structure using Babel tool. The 3-D model of β-lactamase and drug were used for docking. A number of drugs were docked using AutoDock against β-lactamase model. The Lamarckian Genetic Algorithm (LGA) of the Autodock 3.05 was used for docking experiments. Distance-dependent function of the dielectric constant was used for the calculation of the energetic maps and all other parameters were used by default value.

### 2.4 Construction and analysis of phylogenetic tree

The protein sequence of β-lactamase of *A. hydrophila* was used for searching the homology with BLAST and homologous sequences were retrieved Genbank. All these protein sequences were aligned in CLUSTALX 2.0.5. These sequences used passion correction equation which was implemented in the MEGA 4.0 program for construction of a phylogenetic tree by neighbor-joining method. A total of 1000 bootstrapped values were sampled to determine a measure of the support for each node on the consensus tree.

## 3. RESULTS AND DISCUSSION

The size of β-lactamase (NCBI Accession No. ABK35804.1) of *A. hydrophila* subsp hydrophila ATCC 7966 was 253 amino acids long that showed the structural homology with 3-D crystal structure (PDB: 1×8g) of mono zinc carbapenemase (cphA) of *A. hydrophila*. The percentage identity of β-lactamase with template was 86% and positive amino acid residue was 88%. E-value score showed in the BLAST result for selected 3-D structure was 1e-130. In the present study, we observed significant similarity of score and dope score target and template protein. Total five models were generated and selected the one 3-D model β-lactamase which showed similar free energy. The 3-D model of β-lactamase of *A. hydrophila* was shown (Fig. 1). It contains the three major lobe first (red) lobe 2 α-helix and 3 α-helix in green lobe whereas the 9 β-sheets.

**Fig.1.**
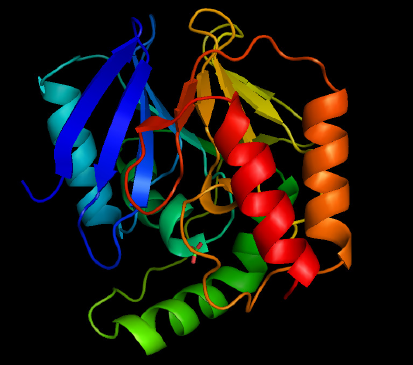
The 3-D model of β-lactamase of *A. hydrophila*.

The similar approach has been previously applied for cry2Ab-type gene of *Bacillus thuringiensis.* The protein model was constructed by homology modeling and predicted a receptor-binding site by docking (Lin et al., 2008). The 3-D models for the 65-kDa of Cry4A and Cry4B endotoxins of *B. thuringiensis* has been constructed by homology modeling based on crystal structure of Cry1Aa and Cry3Aa (Angsuthanasombat et al., 2004). Whereas stereochemical and spatial arrangement of amino acids residues within the most favored region in the Ramachandran plat is determined. The torsion angle of 3-D model of β-lactamase of *A. hydrophila* was 91.1% in favored region whereas 0.9 % amino acid residue in disallowed region. The overall G-factor of 3-D model was 1.0 % as it indicated the best protein model.

In a previous study, homology modeling has been used for generating a 3-D model of β-ketoacyl acyl carrier synthase (KAS) III protein of *Enterococcus faecalis.* The generation of 3-D model has been done by through Modeller that was validated by using PROCHECK programme. In this study, the torsion angle of 88.9 % of residues had within the allowed region and only 0.3 % of residues in disallowed region of the Ramachandran plot. Similar approach has been applied in another study, identification of active site residue using docking and found Naringenin and Apigenin that was able to inhibit the growth of bacteria in culture (Jeong et al., 2007). The β-lactamase is the one of most known enzymes that degrades β-lactam ring. In the present study, total 11 drugs were selected for docking with active amino acid residues of β-lactamase. Out of 11 drugs, 3 drugs (Ampicillin, Astreonam and Sultamicillin) were found to be highest binding energy means lowest docked energy (Table 1). The docking energy of Ampicillin, Astreonam and Sultamicillin were −20.80, −21.05 and −20.88 respectively. On the other hand, these drugs have relatively similar interaction energy with protein.

**Table 1.**
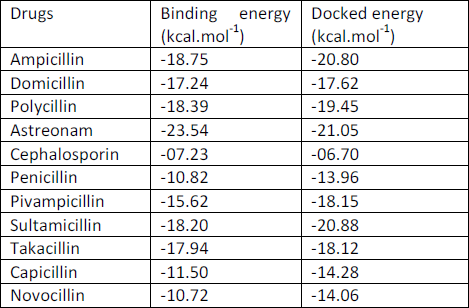
The interaction energy (kcal.mol^-1^) of β-lactamase and drugs obtained by docking.

There is a hydrogen bond (HB) observed that can be helpful for stronger binding between protein and drug molecules. The active amino acid residues of β-lactamase showed the affinity with Ampicillin, Astreonam and Sultamicillin were given (Fig. 2, 3, 4).

**Fig.2.**
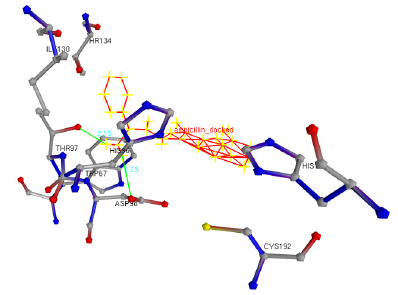
The interaction of high affinity Ampicillin drugs with β-lactamase.

**Fig.3.**
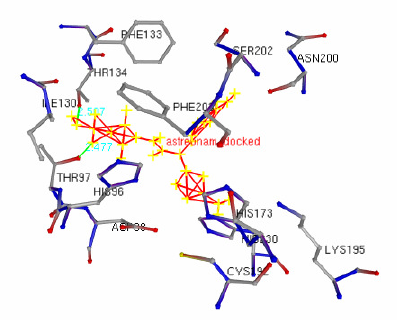
The interaction of high affinity Astreonam drugs with β-lactamase.

**Fig.4.**
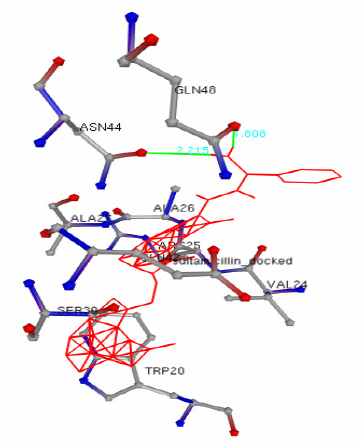
The interaction of high affinity Sultamicillin drugs with β-lactamase.

The HB was formed between β-lactamase and Ampicillin in the active amino acid such as Thr97 and Asp98 with 2.15 and 2.25 Ǻ distances (Fig. 2). In other drugs (Astreonam), HB was formed with active amino acids that include Thr97 and Thr34 with the 2.477 and 2.507Ǻ distances (Fig. 3). While HB was formed between β-lactamase and Sultamicillin such amino acids Gln48 and Asn44 with1.808 and 2.215 Ǻ distances (Fig. 4).

The phylogenetic tree of β-lactamase of *A. hydrophila* and other potential bacteria were constructed. Therefore, homology of β-lactamase was present in a number of bacteria. As depicted in Fig.5, phylogenetic tree of the β- lactamase of *A. hydrophila* with other bacteria that contains homologous protein. The β-lactamase of *A. hydrophila* showed the closely related with *A. sobria, A. salmonicida, A. caveai, A. veronii, A. jandaei* and *A. allosaccharophila* which was present in the same clade.

**Fig.5.**
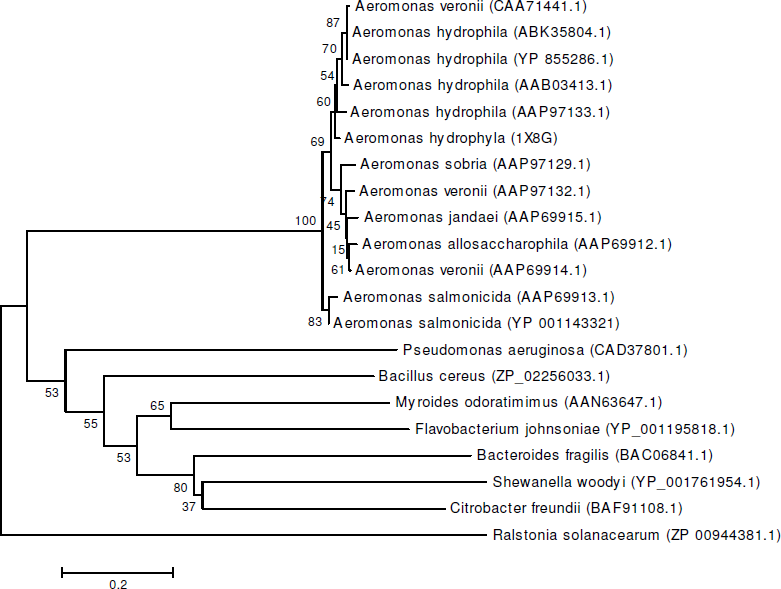
Phylogenetic tree of β-lactamase of *A. hydrophila* with other significant bacteria. The bar 0.2 represent the changes per site.

The other groups of significant bacteria present in other clades.

However, these bacteria contain homologous region of β- lactamase. It indicates that Ampicillin, Astreonam and Sultamicillin can also be used for controlling of growth of *Aeromonas* species and also used for other bacteria. The phylogeny indicates that β-lactamase is a stable target for genetic relationship. In a previous study, nucleotide sequences of gyrB have been used for confirmation of 53 *Aeromonas* strains including some new isolates that was characterized by 16S rDNA sequences (Yanez et al., 2003). The phylogeny of *Aeromonas bestiarum* and *Aeromonas salmonicida* has been determined in 70 strains using rpoD sequence encoding the sigma70 factor. This analysis was complemented with the sequence of gyrB which had already proven useful for determining the phylogenetic relationship.

## CONCLUSION

We have successfully constructed a 3-D model for β- lactamase of *A. hydrophila* that could be used for screening of potent drugs. We found that Ampicillin, Astreonam and Sultamicillin have highest binding affinity. It means that stronger inhibition of growth of *A. hydrophila*. Whereas, phylogeny of β-lactamase of *A. hydrophila* indicates that the same drug can be inhibited the growth of other bacteria. This study also provides a new insight for controlling of unwanted and heavy use of drug in lab practices. The excessive use of antibiotics has accelerated the spreading of drug-resistant strains that may be a grand challenge.

## ACKNOWLEDGEMENT

Authors thank Indra Mani and Satya Prakash for their critical review and suggestion.

